# Aberrant modulation of brain activity underlies impaired working memory following traumatic brain injury

**DOI:** 10.1101/2021.03.23.436708

**Authors:** Abbie S. Taing, Matthew E. Mundy, Jennie L. Ponsford, Gershon Spitz

## Abstract

Impaired working memory capacity is a common and disabling consequence of traumatic brain injury (TBI) that is caused by aberrant neural processing. However, due to high heterogeneity in results across studies, it is challenging to conclude whether impaired working memory in this population is driven by neural hypo- or hyper-activation, and the extent to which deficits are perpetuated by specific working memory subprocesses. Using a combined functional magnetic resonance imaging and working memory paradigm, we tested the hypothesis that the pattern of neural activation subserving working memory following TBI would interact with both task demands and specific working memory subcomponents: encoding, maintenance, and retrieval. Behaviourally, we found that working memory deficits were confined to the high cognitive load trials. Our results confirmed our key prediction. Overall, TBI participants showed reduced brain activity while performing the working memory task. However, interrogation of the subcomponents of working memory revealed a more nuanced pattern of activation. When we simply averaged across all task trials, regardless of cognitive load or subcomponent, TBI participants showed reduced neural activation. When examined more closely, patterns of brain activity following TBI were found to interact with both task demands and working memory subcomponent. Participants with TBI demonstrated an inability to appropriately modulate brain activity between low and high demand conditions necessary during encoding and maintenance stages. Therefore, we demonstrate that conclusions about aberrant neural processing are dependent upon the level of analysis and the extent to which general cognitive domains can be parcellated into its constituent parts.

## Introduction

Traumatic brain injury (TBI) is a debilitating condition that impairs a range of cognitive domains (Draper & Ponsford, 2008; Ponsford et al., 2014). Working memory is often affected, which comprises distinct cognitive subprocesses that integrate, store, and manipulate information online for temporary use (Baddeley, 2003; Baddeley & Hitch, 1974). Due to the inherent utility of these cognitive subprocesses for many of our everyday activities, impaired working memory following TBI significantly disrupts resumption of function (Avery, Smillie, & de Fockert, 2013; Baddeley, 2010; Burgess, Gray, Conway, & Braver, 2011; McVay & Kane, 2012). Despite the relevance of intact working memory for adaptive function, little has been done to understand the role of the specific subprocesses in perpetuating these deficits.

In healthy individuals, working memory is supported by a distributed network involving frontoparietal (Rottschy et al., 2012) and temporal regions (Lee & Rudebeck, 2010; Schon, Quiroz, Hasselmo, & Stern, 2009). These regions are critical for the integration of information necessary for decision-making (De Pisapia, Slomski, & Braver, 2007; C. Kim, Kroger, Calhoun, & Clark, 2015), updating of information (Murty et al., 2011), and coordinating attentional resources (Rossi, Pessoa, Desimone, & Ungerleider, 2009).

The network of regions critical for working memory is also most susceptible to the direct physical forces and secondary pathology of TBI (Bigler, 2001). Functionally, this neural network displays aberrant neural activity following TBI, leading to impaired working memory capacity (Kasahara et al., 2011). This is particularly evident as task complexity and effort increases (Perlstein et al., 2004; Sanchez-Carrion, Fernandez-Espejo, et al., 2008; Sanchez-Carrion, Gomez, et al., 2008). Others, however, have demonstrated aberrant neural processes in this distributed network in the absence of significant working memory deficits (Christodoulou et al., 2001; McAllister et al., 2001), potentially highlighting longer-term restorative, compensatory, or neuroplastic processes (Hillary, 2008).

The field is also mixed with respect to the *direction* of aberrant neural processing during working memory. That is, whether working memory deficits are driven by hypo-(Sanchez-Carrion, Fernandez-Espejo, et al., 2008; Sanchez-Carrion, Gomez, et al., 2008) or hyper-activation (Christodoulou et al., 2001; McAllister et al., 2001; Perlstein et al., 2004), relative to healthy controls. These disparities likely reflect methodological differences across studies that are not taken into account when interpreting the results. One key variable is time since injury (Dunning, Westgate, & Adlam, 2016). For example, TBI participants showed an initial (average 8 months following injury) neural hypoactivation, relative to controls, which progressively resolved 6 months later and coincided with significant improvements in working memory performance (Sanchez-Carrion, Fernandez-Espejo, et al., 2008). Thus, the time at which individuals are assessed can reveal different patterns of aberrant brain activity.

We proposed that another significant factor contributing to inconsistent findings in the field is the precision with which working memory is measured. Although multiple subprocesses are known to support working memory capacity (Fletcher & Henson, 2001; H. Kim, 2019), working memory is generally measured using omnibus tests. For example, the n-back task, which has been used in several functional magnetic resonance imaging (fMRI) studies (McAllister et al., 2001; Perlstein et al., 2004; Sanchez-Carrion, Fernandez-Espejo, et al., 2008; Sanchez-Carrion, Gomez, et al., 2008), does not delineate specific subprocesses since the sequential nature of the task requires execution of these subprocesses simultaneously (Jaeggi, Buschkuehl, Perrig, & Meier, 2010). Studies in healthy individuals have shown the pattern of neural activation differs depending on the subprocesses of working memory (e.g. Narayanan et al., 2005). Despite this, there has been limited investigation of these specific subprocesses in individuals with TBI.

Delayed match-to-sample behavioural paradigms allow measurement and analysis of working memory subcomponents by demarcating stages of encoding, maintenance, and retrieval (Jensen, Gelfand, Kounios, & Lisman, 2002). To our knowledge, only one previous study in paediatric TBI (Newsome et al., 2008) has combined this behavioural paradigm with functional magnetic resonance imaging (fMRI) to investigate the neural basis of working memory disruption in individuals with moderate to severe TBI. This single study highlights two key results: 1) neural hyperactivity, relative to controls, in frontal, temporal, and occipital regions as task demands increased, which was *specific* to encoding and retrieval subcomponents; 2) neural hypoactivation, relative to controls, in prefrontal and parietal regions, which was *specific* to the maintenance subcomponent of working memory. These findings have been interpreted as reflecting reduced capacity to differentially allocate neural resources during the maintenance phase, alongside compensatory over-activation during encoding and retrieval. However, as there are potential differences in pathophysiology, developmental, and psychosocial factors associated with paediatric TBI (Ponsford, Sloan, & Snow, 2012), it remains unclear whether these findings can be generalised to the adult TBI population.

Here, we used a delayed match-to-sample task combined with fMRI to investigate working memory deficits in adults with moderate to severe TBI. To first establish the sensitivity of our behavioural task, and ascertain the presence of chronic working memory deficits, we characterised behavioural performance across both acute and chronic TBI cohorts relative to healthy controls. We subsequently investigated *whether* and *how* observed behavioural deficits were associated with neural changes during specific working memory subcomponents using fMRI. Based on previous key studies (Perlstein et al., 2004; Sanchez-Carrion, Fernandez-Espejo, et al., 2008; Sanchez-Carrion, Gomez, et al., 2008), we hypothesised that participants with TBI would display working memory deficits that were conditional on task demands (i.e. greater deficits evident with increased task demands). Furthermore, based on the single previous study in adolescents (Newsome et al., 2008), we hypothesised that the *direction* of neural activation would depend on the working memory subcomponent: relative to healthy individuals, the TBI group were predicted to show increased activation during encoding and retrieval but decreased activation during maintenance.

## Participants and Methods

The study was approved by Monash Health/University Human Research Ethics Committee and conducted in accordance with the Declaration of Helsinki. Written informed consent was obtained from all participants. Participants were recruited from the TBI rehabilitation program of Epworth Healthcare, either from successive admissions to the inpatient ward or, for chronic participants, via a longitudinal follow-up database. Exclusion criteria included age < 18 or > 75 years, prior history of TBI or other neurological conditions, significant psychiatric or substance abuse history, and MRI contraindication. Forty-three individuals with moderate to severe TBI, as determined using the Westmead Post Traumatic Amnesia Scale (WPTAS), participated in the study (see Table 1 and Supplementary Table 1 for participant details; see Supplementary Fig. 1 for TBI lesion overlay map). Of these, 25 were inpatients (17 males, 8 females) in the acute phase of recovery (*M =* 2.16 months, *SD* = 1.48 months, range = 0.69 – 6.64 months) and 18 individuals (14 males, 4 females) were in the chronic phase of recovery (*M* = 23.44 months, *SD* = 6.76 months, range = 13.35 – 34.82 months). Thirty-eight healthy controls (26 males, 12 females) of similar age, sex, and education were also recruited. There were no significant differences between the TBI groups and healthy controls on any of the demographic variables; nor were any significant difference between the two TBI groups on injury factors (all comparisons *P* > 0.05). All participants contributed to our first aim, characterising working memory behavioural performance. However, only a subset of participants completed the fMRI working memory task (18 chronic TBI and 17 healthy controls), thus contributing to our second aim of linking neural activity with behaviour. Three participants (2 acute TBI and 1 healthy control) were excluded from the behavioural analysis due to technical errors with the response button and apparent poor effort on task (i.e. no variation in response).

**Table 1.** Demographic and clinical information of participants

| **Demographic variables** | **Acute TBI,**<br>***M (SD)*** | **Chronic TBI,**<br>***M (SD)*** | **Healthy controls,**<br>***M (SD)*** |
| --- | --- | --- | --- |
| Age (years) | 38.36 (16.82) | 44.11 (15.79) | 40.05 (17.14) |
| Gender (male/female) | 17/8 | 14/4 | 26/12 |
| Education (years) | 13.82 (3.27) | 14.81 (2.40) | 14.68 (2.79) |
| Time since injury (months) | 2.16 (1.51) | 23.44 (6.96) | - |
| PTA (days)<sup>a</sup> | 22.46 (14.63) | 33.12 (38.88) | - |
| GCS (lowest)<sup>b</sup> | 9 (4.37) | 9.47 (4.14) | - |

GSC = Glasgow Coma Scale; PTA = post-traumatic amnesia.
<sup>a</sup> PTA duration were available for $n = 41$ TBI participants.
<sup>b</sup> Acute GCS were available for $n = 42$ TBI participants.

### Working Memory Paradigm

Working memory was examined using the Sternberg delayed match-to-sample task (Fig. 1; Sternberg, 1966). In this task, participants are required to remember a memory set consisting of items (letters) presented on a screen, which they are asked to encode into working memory (encoding). After a short delay (maintenance), a single item is presented, and participants respond by indicating whether the item was in the previous memory set (retrieval). To modulate cognitive load/task demands, participants were required to encode either a two-item (low cognitive load) or six-item (high cognitive load) memory set. This task consisted of 28 blocks, alternating between low and high cognitive load conditions. Each block comprised 14 trials. Items were presented for 0.8 seconds followed by a 0.2 second inter-stimulus interval. The duration of encoding was 1.8 seconds for the low cognitive load condition and 5.9 seconds for the high cognitive load condition. The duration of maintenance was 5.2 seconds across both load conditions. The duration of retrieval was modelled as the participant’s reaction times.

**Figure 1.**
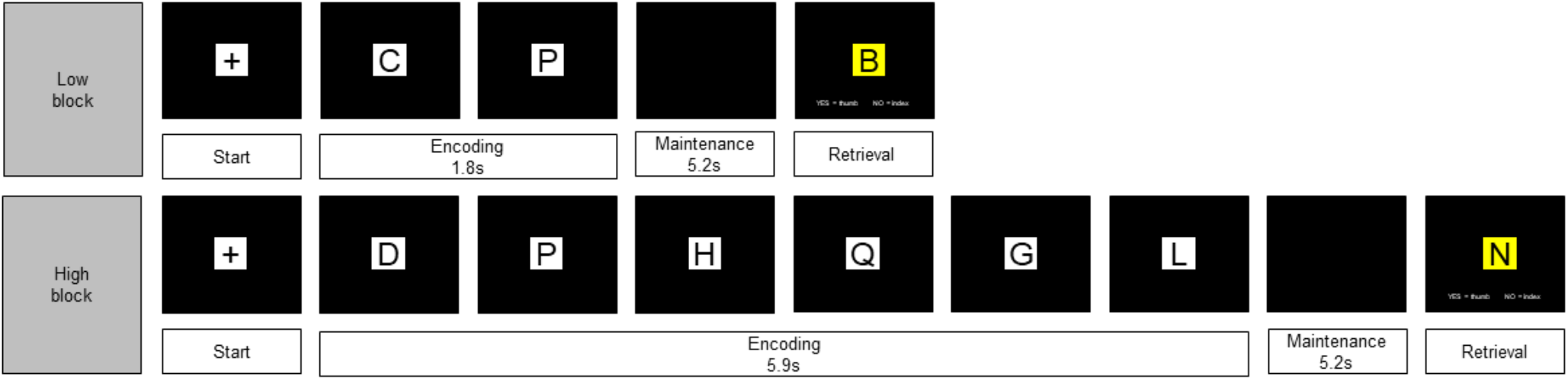
Schematic diagram of low and high cognitive load trials of the Sternberg working memory task. Two (low cognitive load) or six (high cognitive load) letters were presented during the encoding phase, followed by a maintenance phase in which the screen was blank, and finally a probe was presented during retrieval phase. Low and high cognitive load conditions were presented alternately.

### MRI Acquisition

Structural and functional MR images were acquired with a 3.0 Tesla Siemens Magnetom Skyra scanner (Monash Biomedical Imaging, Clayton, Australia) and 32-channel head coil. Functional images were obtained using single-shot gradient-echo planar imaging (EPI) with the following parameters: repetition time (TR) = 0.74 seconds; echo time (TE) = 39 milliseconds; simultaneous multi-slice (SMS) acceleration factor = 8; flip angle = 52°; 210 × 210 matrix; voxel size = 2.4 × 2.4 × 2.4 mm. A single-band reference scan was also obtained for EPI registration purposes with the following parameters: TR = 6.37 seconds; TE = 39 milliseconds; flip angle = 52°; 210 × 210 matrix; voxel size = 2.4 × 2.4 × 2.4 mm. To correct for B0 inhomogeneity in the EPI scans, a field map was acquired using a double-echo spoiled gradient echo sequence with the following parameters: TR = 0.68 seconds; TE = 4.92/7.38 milliseconds, flip angle = 60°, 210 x 210 matrix, 3.3 x 3.3 x 2.4 mm. A high-resolution 3D T1-weighted image covering the entire brain was also acquired with the following parameters: TR = 2.0 seconds; TE = 2.03 milliseconds; flip angle = 8°; ascending order; 256 × 256 matrix; voxel size = 1.0 × 1.0 × 1.0 mm.

### Statistical Analysis

#### Behavioural and Demographic Data

Behavioural and demographic data were analysed using R version 3.6.0 (R Core Team, 2019). Independent samples t-tests were used to assess group differences on the demographic variables including age, sex, and years of education. Behavioural data were screened and assessed for violation of statistical assumptions prior to analysis. Our behavioural measures of interest were accuracy and reaction time on the working memory task. Accuracy was determined using dprime, a sensitive index which measures an individual’s ability discriminate signal from noise. Reaction times were calculated using the average reaction time per load. Age and education were added as covariates in all models given they could affect working memory performance (Fitzpatrick, Archambault, Janosz, & Pagani, 2015; Mattay et al., 2006) and reaction time (Der & Deary, 2017; Tun & Lachman, 2008). Linear mixed models were used to analyse working memory accuracy and reaction time to better account for clustering and non-independence of measures within participants. In the first model, we used a linear mixed model to model group and load as fixed effects, and participant as a random effect. In a second model, we subsequently added a *group × load* interaction term to investigate whether cognitive load/task demands moderate accuracy and reaction time on the task. Given our a priori hypothesis of a specific group difference for high cognitive load trials, we conducted linear regressions for each load condition (i.e. low and high) separately. Post-hoc analyses were followed up using two-tailed independent samples t-tests with multiple comparison correction.

#### Imaging Data

##### MRI Preprocessing

Lesions were manually segmented using MRIcron (http://www.mricro.com/mricron) and subsequently preprocessed using fMRIPrep 20.0.0 (Esteban et al., 2018). The following steps were applied: undistortion of EPI data, realignment, normalisation, and estimation of confounds. Further information about the MRI preprocessing can be found in the Supplementary (see “Detailed MRI Preprocessing”).

##### fMRI Analysis

Imaging data were analysed using FSL FEAT version 6.0.2 (FMRIB’s Software Library, www.fmrib.ox.ac.uk/fsl). Onset times for encoding were modelled as the first stimulus presentation of each load condition; low cognitive load trials were modelled as 1.8 seconds and high cognitive load trials modelled as 5.9 seconds in length. Onset times for maintenance were modelled as the period immediately following the last stimulus presentation of each load; the maintenance period duration was 5.2 regardless of load condition. Onset times for retrieval was modelled as the length of time of the probe stimulus presentation; this was equivalent to the participant’s reaction/response times on the trial. In the first level FEAT model, we modelled the main effect of load (i.e. low and high), subcomponent (i.e. encoding, maintenance, and retrieval) and their interaction for each participant. To reduce motion-related artifact, additional anatomical CompCor regressions were also added at the first level (Muschelli et al., 2014). We investigated group-level differences using FLAME 1+2 mixed effects with automatic outlier de-weighting. Imaging findings are reported using a cluster level threshold of *Z* > 3.1 and a family wise error cluster correction of *P* < 0.05.

## Results

### Working memory deficits in TBI are conditional on task demands

We used mixed linear models to investigate the association between group (healthy controls vs acute TBI; healthy controls vs chronic TBI), load (low vs high), and working memory performance (accuracy and reaction time). Overall, TBI participants were significantly less accurate at both *acute* (*t*(148) = −2.37, 95% CI [-0.66 – −0.06], *P* = 0.018) and *chronic* timepoints (*t*(148) = −1.97, 95% CI [-0.64 – 0.001], *P* = 0.049). *Acute* TBI participants were significantly slower to respond during trials, compared to healthy controls (*t*(148) = 3.89, 95% CI [0.13 – 0.40], *P* < 0.001). No reaction time difference was apparent between *chronic* TBI participants and healthy controls (*t*(148) = −0.50, 95% CI [-0.18 – 0.11], *P* = 0.618). As expected, task accuracy was significantly higher in low cognitive load condition, compared to high cognitive load condition, regardless of group (*t*(148) = −13.37, 95% CI [-1.27 – −0.94], *P* < 0.001). Reaction time was significantly faster in the low cognitive load condition, compared to the high cognitive load (*t*(148) = 9.60, 95% CI [0.20 – 0.31], *P* < 0.001).

To examine whether cognitive load moderated the relationship between group and task performance (accuracy and reaction time), we subsequently added a *group* × *load* interaction into the model. Cognitive load did not moderate the relationship between group and task accuracy for the *acute* TBI cohort, though this was trending towards significance (*t*(146) = − 1.83, 95% CI [-0.72 – 0.03], *P* = 0.068). Cognitive load did not moderate the relationship between group and task accuracy for the *chronic* TBI cohort (*t*(146) = −1.27, 95% CI [-0.66 – 0.14], *P* = 0.204). Similarly, cognitive load did not moderate the relationship between group and reaction time for both *acute* (*t*(146) = −0.79, 95% CI [-0.17 – 0.07], *P* = 0.428) and *chronic* TBI cohorts (*t*(146) = −0.36, 95% CI [-0.16 – 0.11], *P* = 0.722).

Despite not finding a significant interaction effect in using linear mixed models, we further probed the relationship between load and group using linear regression due to our a priori hypotheses. In line with our predicted hypothesis, task accuracy did not differ between groups in the low load condition (*acute* TBI vs healthy controls: *t*(73) = −1.21, 95% CI [-0.52 – 0.13], *P* = 0.232; *chronic* TBI vs healthy controls: *t*(73) = −1.15, 95% CI [-0.55 – 0.15], *P* = 0.252; Fig. 2A). However, TBI participants were significantly less accurate in the high load condition at both *acute* (*t*(73) = −2.57, 95% CI [-0.93 – −0.12], *P* = 0.012) and *chronic* timepoints (*t*(73) = −2.07, 95% CI [-0.88 – 0.004], *P* = 0.048; Fig. 2B). *Acute* TBI participants were significantly slower to respond, compared to healthy controls, during both the low (*t*(73) = −3.73, 95% CI [0.13 – 0.44], *P* < 0.001; Fig. 2C) and high cognitive load conditions (*t*(73) = 3.16, 95% CI [0.09 – 0.39], *P* = 0.002; Fig. 2D). However, no reaction time differences were apparent between *chronic* TBI participants and healthy controls in both the low (*t*(73) = −0.23, 95% CI, −0.19 – 0.15, *P* = 0.821; Fig. 2C) and high cognitive load conditions (*t*(73) = −0.66, 95% CI, − 0.22 – 0.11, *P* = 0.513; Fig. 2D).

**Figure 2.**
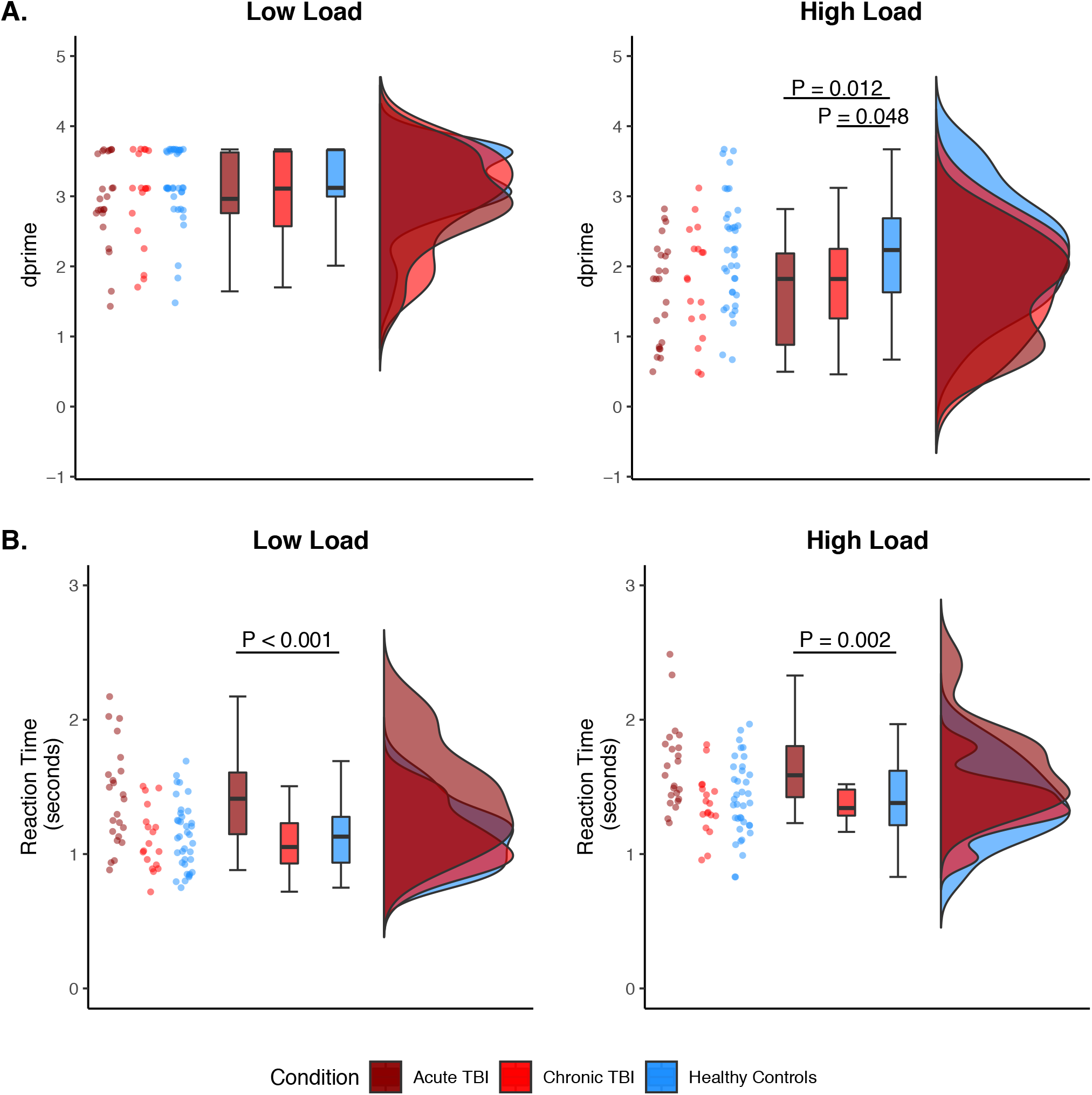
Behavioural results for the Sternberg working memory task. A) Plots depicting accuracy, as measured using dprime (higher values denote better performance). Individual datapoints are displayed along with violin plots and boxplots showing distribution. Performances were comparable across groups in the low cognitive load condition (*P >* 0.05; panel A, left). In the high cognitive load condition, however, performance was significantly impaired for both the acute (*P* = 0.012) and chronic TBI groups (*P* = 0.048; panel A, right). B) Plots representing reaction time (higher values denote slower performance). The acute TBI group was significantly slower than healthy controls in both the low (*P* < 0.001; panel B, left) and high cognitive load condition (*P =* 0.002; panel B, right). There were no significant differences in reaction time between the chronic TBI group and healthy controls in both load conditions (*P >* 0.05; panel B, left and right). Note: *P*-values were adjusted for multiple comparisons.

### Sternberg task activates the expected brain network associated with working memory

Prior to testing our key fMRI analyses, we first assessed whether our Sternberg memory task activated the expected ‘working memory’ brain network in our study sample (i.e. 18 chronic TBI participants and 17 healthy controls; Fig. 3A and Supplementary Table 2). There were areas of overlap between the various subcomponents in frontal and occipital areas, the insula and cerebellum. Encoding was associated with increased activation in temporal areas (e.g. left middle temporal gyrus, right superior temporal gyrus). Retrieval was associated with increased activation in the left angular gyrus. In general, these results overlap with meta-analytic working memory mask derived from neurosynth (https://www.neurosynth.org/;Fig. 3B). However, consistent with the visual nature of our task, we found more activation in occipital areas compared to the meta-analytic mask (which includes results from working memory studies of various modalities).

**Figure 3.**
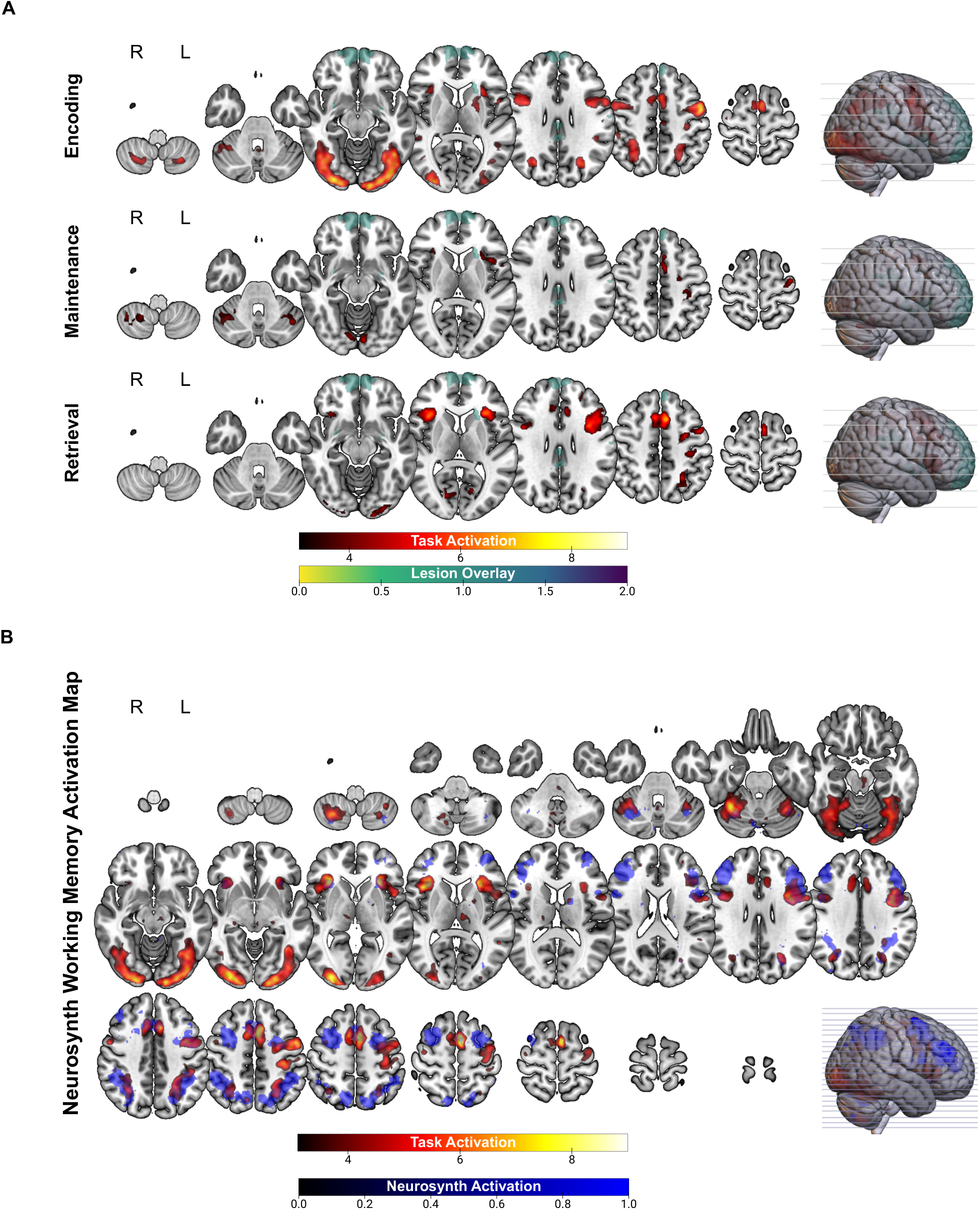
Activation during the Sternberg working memory task for the fMRI cohort (18 chronic TBI, 17 healthy controls). A) Pattern of activation segregated according to encoding, maintenance, and retrieval stages (red/yellow). There was minimal overlap between these clusters and TBI lesions (green). All subcomponents were associated with increased activation in frontal and occipital areas, the insula and cerebellum. Encoding was associated with increased activation in temporal areas (e.g. left middle temporal gyrus, right superior temporal gyrus). Retrieval was associated with increased activation in the left angular gyrus. B) Overall activation during the Sternberg working memory task (red/yellow) overlaid on a working memory meta-analytic mask (blue) derived from neurosynth.

**Figure 4.**
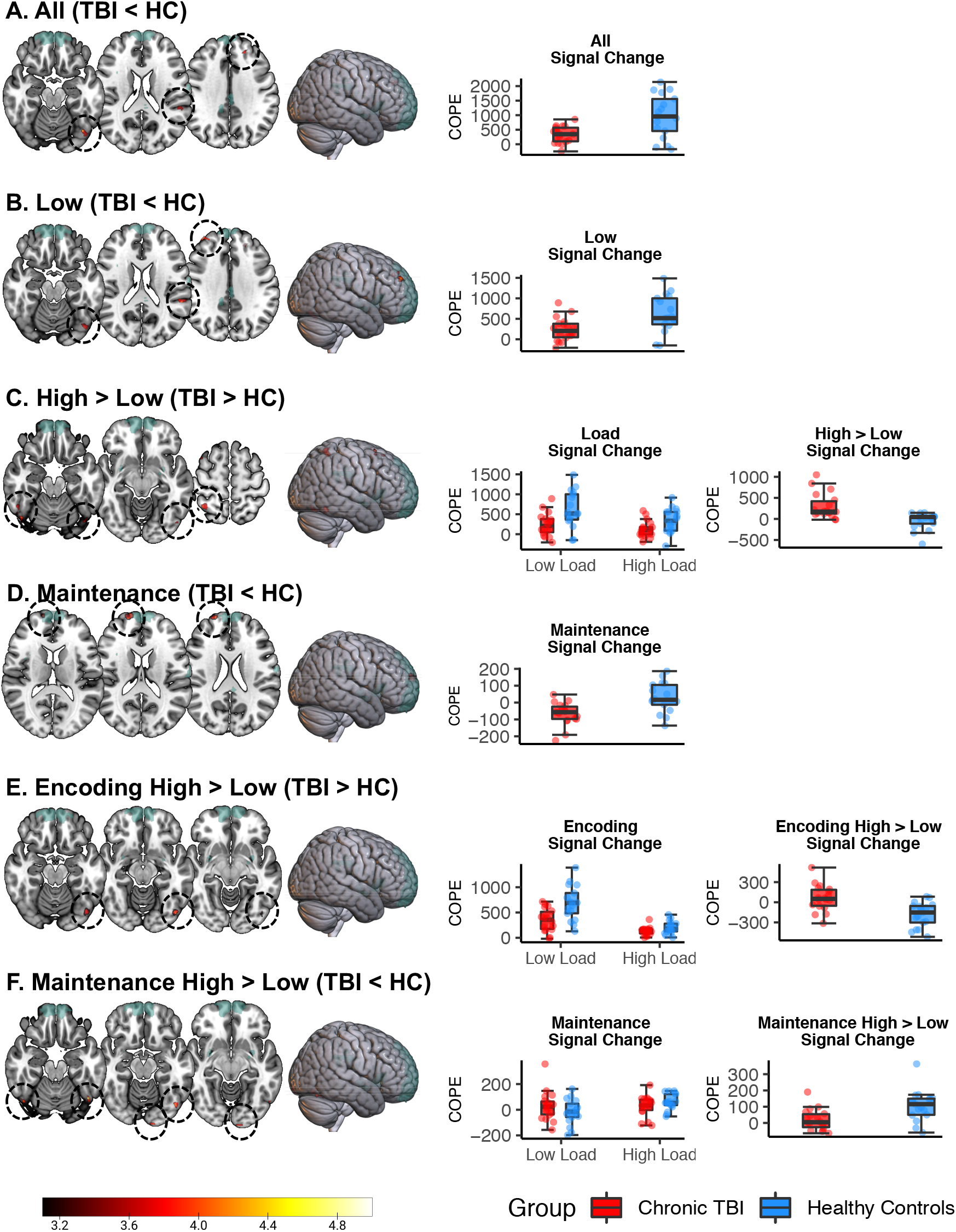
fMRI group differences on the Sternberg working memory task. There was minimal overlap between significant clusters (red/yellow) and TBI lesions (green). A) Average across all trials (i.e. regardless load and subcomponent), the TBI group showed reduced activation in the left cerebellum, left middle frontal gyrus, and left superior temporal gyrus. Panel A right, boxplot overlaid with individual datapoints comparing signal change when averaged across all trials between the TBI group and healthy controls. B) The TBI group showed reduced activation in the low cognitive load condition (left cerebellum, right middle frontal gyrus, left superior temporal gyrus, and left superior frontal gyrus). Panel B right, comparison of signal change during the low cognitive load condition between the TBI group and healthy controls. C) With increased cognitive load, the TBI group showed attenuated reduction in activity in the right superior parietal gyrus, right inferior temporal gyrus, right middle frontal gyrus, and left inferior occipital gyrus). Panel C middle, comparison of signal change in the low and high cognitive load conditions between the TBI group and healthy controls. Panel C right, comparison of signal change with increased cognitive load (i.e. high > low) between the TBI group and healthy controls. D) During maintenance, the TBI group showed reduced activation in the right dorsolateral prefrontal cortex. Panel D right, comparison of signal change during maintenance between the TBI group and healthy controls. E) With increased cognitive load, the TBI group showed increased activation during encoding in the left inferior occipital gyrus. Panel E middle, comparison of signal change during encoding of low and high load conditions between the TBI group and healthy controls. Panel E right, comparison of signal change during encoding with increased cognitive load (i.e. encoding high > encoding low) between the TBI group and healthy controls. F) In contrast, the TBI group displayed decreased activation during maintenance in bilateral cerebellum and left calcarine sulcus. Panel F middle, comparison of signal change during maintenance of low and high conditions between the TBI group and healthy controls. Panel F right, comparison of signal change during maintenance with increased cognitive load (i.e. maintenance high > maintenance low) between the TBI group and healthy controls.

### Working memory following TBI is characterised by reduced neural activation when disregarding cognitive load and working memory subcomponent

The TBI group showed reduced activation in comparison to healthy controls when neural activation was averaged across all trials, regardless of load or working memory subcomponent. This ‘hypoactivation’ was present in three clusters: left cerebellum, left middle frontal gyrus, and left superior temporal gyrus (Fig. 4A and Supplementary Table 3). Healthy controls did not show reduced patterns of activation in comparison to the TBI group.

### Individuals with TBI show aberrant modulation of brain activity with increased task demands

During low cognitive load trials, TBI participants showed reduced activity compared to healthy controls in several brain regions. This ‘hypoactivation’ was found in the left cerebellum, right middle frontal gyrus, left superior temporal gyrus, and left superior frontal gyrus (Fig. 4B and Supplementary Table 4). Interestingly, no group difference was found for high cognitive load trials. However, relative to controls, TBI participants showed attenuated reduction in activity with increased working memory load in four brain regions: right superior parietal gyrus, right inferior temporal gyrus, right middle frontal gyrus, and left inferior occipital gyrus (Fig. 4C and Supplementary Table 5).

### Patterns of aberrant brain activation following TBI interact with working memory subcomponent and task demand

As predicted, the *direction* of aberrant brain activation following TBI depended on the working memory subcomponent and cognitive load. TBI participants showed ‘hypoactivation’ during working memory maintenance, relative to healthy controls, regardless of load condition. This reduction in activation was present in the right dorsolateral prefrontal cortex (Fig. 4D and Supplementary Table 6). However, no significant group differences were apparent during working memory encoding or retrieval.

In addition, activation of working memory subcomponents differed as cognitive demands increased. As cognitive load increased, the TBI group showed relatively greater activation in the left inferior occipital gyrus, that was specific to the encoding subcomponent (Fig. 4E and Supplementary Table 7). In addition, with increased task demands, the TBI group showed relatively reduced activation during maintenance, relative to controls, in two clusters: bilateral cerebellum and left calcarine sulcus (Fig. 4F and Supplementary Table 8).

## Discussion

This study tested the hypothesis that changes in working memory following TBI would be dependent on task demands and specific working memory subcomponents. We used a combined fMRI and a delayed match-to-sample behavioural task to probe working memory encoding, maintenance, and retrieval. In doing so, we identified that changes in working memory capacity following TBI are underpinned by aberrant neural processing that manifests conditionally depending on task demands and specific working memory subcomponents. Behaviourally, we showed that individuals with TBI displayed impaired working memory deficits that were confined to the high cognitive load trials, at both acute and chronic timepoints following TBI. Neurally, we showed that aberrant brain activation during working memory may be characterised as hypo- or hyper-activation, depending on both task demands and the working memory subcomponent. TBI participants demonstrated increased activation in the left occipital gyrus during encoding whereas decreased activation was observed in the bilateral cerebellum and left calcarine sulcus during maintenance. The present results suggest that individuals with TBI have impaired capacity to appropriately modulate brain activity with increased task demands specific to encoding and maintenance stages.

Our behavioural results are consistent with past studies that have observed greater difficulties with working memory as task demands increased (Perlstein et al., 2004; Sanchez-Carrion, Fernandez-Espejo, et al., 2008; Sanchez-Carrion, Gomez, et al., 2008). In the current study, working memory impairments were confined to the high cognitive load trials for both acute and chronic TBI participants, indicating the persistence of working memory deficits after TBI. Interestingly, we found that reaction time findings varied according to recovery phase. That is, acute TBI participants were significantly slower in both load conditions whereas chronic TBI participants performed comparably to healthy controls on reaction time regardless of load conditions. One possibility is that slower reaction times may represent generalised impairment in processing speed during early recovery that is unrelated to the presentation of working memory difficulties (Felmingham, Baguley, & Green, 2004). Alternatively, slowed processing speed may contribute to working memory deficits initially whilst other mechanisms may be responsible for maintaining working memory dysfunction in the longer term.

The neural underpinnings of working memory have been previously explored in TBI (Perlstein et al., 2004; Sanchez-Carrion, Fernandez-Espejo, et al., 2008; Sanchez-Carrion, Gomez, et al., 2008). Our results align with these prior studies by demonstrating that impaired working memory following TBI involves aberrant activation in a distributed network of brain regions. Evaluation of task load revealed brain activation with increased task demands following TBI, relative to healthy controls. Similar findings have been reported in other clinical populations including schizophrenia (Guerrero-Pedraza et al., 2012; Pomarol-Clotet et al., 2008) and mild cognitive impairments (Migo et al., 2015). In contrast, studies in healthy individuals have shown reduced brain activation with increased working memory demands, particularly in regions of the default mode network (DMN; Chee & Choo, 2004; McKiernan, Kaufman, Kucera-Thompson, & Binder, 2003; Pyka et al., 2009; Tomasi, Ernst, Caparelli, & Chang, 2006). The DMN is a network that demonstrates increased activation during rest but disengages during task performance (Raichle et al., 2001). Indeed, greater suppression of the DMN has been found to predict better working memory performance (Sambataro et al., 2010). This finding has often been interpreted as reallocation of limited cognitive resources of irrelevant processes to task-relevant processes during task performance (Mayer, Roebroeck, Maurer, & Linden, 2010). Key structures of the DMN include the medial prefrontal cortex, posterior cingulate cortex/precuneus, and lateral parietal and temporal cortices (Mak et al., 2017; Raichle et al., 2001). The middle frontal gyrus has also been implicated (Demertzi et al., 2011; McGeown, Mazzoni, Venneri, & Kirsch, 2009), albeit less consistently. The regions implicated in the present study that showed increased activation with greater working memory load (i.e. right superior parietal gyrus, right inferior temporal gyrus, and right middle frontal gyrus) can be considered broadly falling within the DMN. Thus, greater neural activity of these network of regions as working memory demands increased – evident in the TBI group – suggest an impaired capacity to effectively reallocate neural resources with this increased task demand. Healthy controls, on the other hand, were better able to suppress DMN activity and in turn more efficiency reallocate resources to aid task performance.

We demonstrated that patterns of brain activity following TBI not only depends on task demands, but are also specific to working memory subprocesses. In line with our predicted hypothesis, the TBI group showed increased activation in the left inferior occipital gyrus with increased cognitive load during encoding. Interestingly, the inferior occipital gyrus has rarely been implicated in working memory, but rather in processing of faces (Jacques et al., 2019) as well as emotionally relevant stimuli (Geday, Gjedde, Boldsen, & Kupers, 2003). Despite this, similar findings were reported by Newsome et al. (2008) who found that adolescents with TBI had increased activation in occipital lobe regions (i.e. cuneus, middle occipital gyrus, and lingual gyrus). As interpreted by Newsome et al. (2008), increased activation in these regions may represent a compensatory response for diminished connectivity with key structures (e.g. frontoparietal structures) involved with working memory. However, in addition to occipital regions, Newsome et al. also reported aberrant activation in frontal, temporal, and parietal regions, which did not differentiate between the TBI group and healthy controls in the present study. This indicates that TBI participants are recruiting similar areas during encoding but require additional activation to support working memory function. Thus, greater recruitment of the inferior occipital gyrus may reflect brain organisation, but the extent to which this is adaptative is unclear.

In contrast, participants with TBI demonstrated reduced activation of the dorsolateral prefrontal cortex during working memory maintenance. The dorsolateral prefrontal cortex has been established as a critical structure involved in executive aspects of working memory, including maintenance and manipulation of information (D’Esposito, Postle, & Rypma, 2000) and strategic control (Bor, Cumming, Scott, & Owen, 2004; Bor, Duncan, Wiseman, & Owen, 2003). Furthermore, this region is actively engaged during retention intervals of delayed response tasks (Curtis & D’Esposito, 2003). Reduced activation in this structure therefore suggests that individuals with TBI may have general strategic difficulty in maintaining task representation. Interestingly, we also found that as task demands increased for working memory maintenance, TBI participants demonstrated reduced activation in the left calcarine sulcus and bilateral cerebellum maintenance, compared to healthy controls. The calcarine sulcus has been implicated in maintaining persistent visual representations observed during short-term maintenance (Tallon-Baudry, Bertrand, & Fischer, 2001). Therefore, reduced activation apparent for the TBI group suggests impaired ability to hold visual representation in mind. With increased task demands, increased activity in this region is even more critical since more information must be temporarily maintained in memory. In addition, accumulating evidence suggests the cerebellum supports higher cognitive functions (Hayter, Langdon, & Ramnani, 2007; Ramnani, 2006), including working memory (Emch, von Bastian, & Koch, 2019; Owen, McMillan, Laird, & Bullmore, 2005; Tomlinson, Davis, Morgan, & Bracewell, 2014). The cerebellum has closed-loop projections to cortical regions including areas of the prefrontal cortex via the cortico-cerebellar circuit (Ramnani, 2006). The role of the cerebellum in working memory specifically appears to be related to articulatory or subvocal rehearsal (Ben-Yehudah, Guediche, & Fiez, 2007; Emch et al., 2019). This indicates that TBI participants, in comparison with healthy controls, used fewer rehearsal strategies to help retain information in mind as working memory demands increased. This further supports the interpretation that individuals with TBI have impaired strategy use.

Although our findings are largely consistent with those of Newsome et al. (2008), one key distinction was that we failed to find differences in activation between the TBI group and healthy controls during retrieval. One reason for this difference may be the slight variations in tasks used in the two studies. The task used by Newsome et al. (2008) differed in that stimuli were presented simultaneously during encoding and for shorter durations (1.7 seconds for both low and high load conditions). It is possible that participants in this previous study found the task more difficult, consequently influence uncertainty during working memory retrieval. An alternative explanation may be that neural processing during working memory differs between children and adults. Previous work in typically developing children and adolescents has found similar, albeit more widespread, patterns of neural activation compared to adults (Klingberg, Forssberg, & Westerberg, 2002; Scherf, Sweeney, & Luna, 2006). The more extensive recruitment is thought to reflect neural inefficiencies in developing brain networks (Bathelt, Gathercole, Johnson, & Astle, 2018). Thus, while aberrant neural activation during retrieval may manifest in paediatric TBI, our results suggest this is not the case for adults.

Our study has important implications. By characterising working memory performance across different recovery periods, we showed that deficits manifest at both acute and chronic recovery timepoints. As working memory function is critical to support adaptive functioning, this further highlights the importance of ongoing rehabilitation to optimise working memory function following TBI. We also replicated past studies by showing that these behavioural deficits are associated with aberrant neural processing. However, here we also extended the field by demonstrating that deficits manifest due to disruption of specific working memory subcomponents (i.e. encoding and maintenance stages). This points to the potential for interventions aiming to improve working memory, for example, by providing specific strategies to improve efficiency of encoding or by extending memory representation in mind. Lastly, we demonstrated that conclusions about the neural underpinning of impaired working memory may vary as a function of level of analysis and the extent to which general cognitive domains are parcellated into finer subprocesses. In the current study, we showed that systematic interrogation of task demands and working memory subcomponents is necessary to unpack the patterns of findings and discern whether neural changes are driven by hypo- or hyper-activation. As an example, we may have been unable to uncover differences during encoding if load effects were not considered. More generally, our findings have important theoretical and clinical implications for the relevant wider field with respect to how we use and interpret our measures for diagnosis and prognosis of cognitive impairment.

Despite these important implications, our study findings should be considered in the context of various limitations. One potential confound related to the task is that duration times differed for the various load conditions and subcomponents. It is possible, for example, that another reason why we failed to find significant group differences during the retrieval phase is because it was shorter in duration. However, the fact that we found significant differences between TBI participants and healthy controls during the low encoding load condition despite it also having a short duration suggest that this was not the case. Another limitation was that acute and chronic TBI participants were not matched. For example, compared to the acute cohort, the chronic TBI group were older, slightly more educated, and had longer duration of PTA. Despite this, it is reassuring that both TBI groups cohorts did not significantly differ overall with respect to demographic and injury factors. In the context of these limitations, future research may consider conducting a longitudinal study to investigate the association between behaviour and neural activation. This approach could provide further insights by clarifying whether the neural mechanisms underpinning working memory evolve over time, what neural reorganisation takes place at different recovery timepoints, and whether changes in behavioural performance are associated with specific neural patterns.

In conclusion, our findings indicated impaired working memory performance following TBI that were confined to the high cognitive load condition. Moreover, our key fMRI finding showed that TBI participants displayed patterns of neural activation that interacted with both task demands and working memory subcomponents: increased activation was observed during encoding whereas decreased activation was apparent during maintenance. Taken together, findings indicate an inability to appropriately modulate brain activity according to task demand that is specific to working memory encoding and maintenance.

## Acknowledgements

This work was supported by the Multi-modal Australian ScienceS Imaging and Visualisation Environment (MASSIVE) HPC facility (www.massive.org.au). The authors would like to thank Olivia McConchie at the Monash Epworth Rehabilitation Research Centre and the staff at the Acquired Brain Injury Ward at Epworth Hospital (Richmond) for their help with recruitment. The authors would also like to thank the staff at Bridge Road Imaging and Monash Biomedical Imaging for their assistance with data collection. Lastly, the authors would like to thank the participants who participated in the study.

## Abbreviations

TBI: traumatic brain injury
fMRI: functional magnetic resonance imaging
WPTAS: Westmead Post Traumatic Amnesia Scale
GCS: Glasgow Coma Scale
EPI: echo planar imaging
TR: repetition time
TE: echo time
SMS: simultaneous multi-slice
COPE: contrast of parameter estimates
BOLD: blood oxygenation level-dependent
DMN: default mode network

## References

Avery, R. E., Smillie, L. D., & de Fockert, J. W. (2013). The role of working memory in achievement goal pursuit. Acta psychologica, 144(2), 361–372.

Baddeley, A. (2003). Working memory: looking back and looking forward. Nat Rev Neurosci, 4(10), 829–839. doi:10.1038/nrn1201

Baddeley, A. (2010). Working memory. Current Biology, 20(4), R136–R140.

Baddeley, A., & Hitch, G. (1974). Working memory. Psychology of learning and motivation, 8, 47–89.

Bathelt, J., Gathercole, S. E., Johnson, A., & Astle, D. E. (2018). Differences in brain morphology and working memory capacity across childhood. Developmental Science, 21(3), e12579.

Ben-Yehudah, G., Guediche, S., & Fiez, J. A. (2007). Cerebellar contributions to verbal working memory: beyond cognitive theory. The Cerebellum, 6(3), 193–201.

Bigler, E. D. (2001). The lesion (s) in traumatic brain injury: Implications for clinical neuropsychology. Archives of clinical neuropsychology, 16(2), 95–131.

Bor, D., Cumming, N., Scott, C. E., & Owen, A. M. (2004). Prefrontal cortical involvement in verbal encoding strategies. European Journal of Neuroscience, 19(12), 3365–3370.

Bor, D., Duncan, J., Wiseman, R. J., & Owen, A. M. (2003). Encoding strategies dissociate prefrontal activity from working memory demand. Neuron, 37(2), 361–367.

Burgess, G. C., Gray, J. R., Conway, A. R., & Braver, T. S. (2011). Neural mechanisms of interference control underlie the relationship between fluid intelligence and working memory span. Journal of experimental psychology: general, 140(4), 674.

Chee, M. W., & Choo, W. C. (2004). Functional imaging of working memory after 24 hr of total sleep deprivation. Journal of Neuroscience, 24(19), 4560–4567.

Christodoulou, C., DeLuca, J., Ricker, J., Madigan, N., Bly, B., Lange, G., … Diamond, B. (2001). Functional magnetic resonance imaging of working memory impairment after traumatic brain injury. Journal of Neurology, Neurosurgery & Psychiatry, 71(2), 161–168.

De Pisapia, N., Slomski, J. A., & Braver, T. S. (2007). Functional specializations in lateral prefrontal cortex associated with the integration and segregation of information in working memory. Cerebral cortex, 17(5), 993–1006.

Demertzi, A., Soddu, A., Faymonville, M.-E., Bahri, M. A., Gosseries, O., Vanhaudenhuyse, A., … Luxen, A. (2011). Hypnotic modulation of resting state fMRI default mode and extrinsic network connectivity. Progress in brain research, 193, 309–322.

Der, G., & Deary, I. J. (2017). The relationship between intelligence and reaction time varies with age: Results from three representative narrow-age age cohorts at 30, 50 and 69 years. Intelligence, 64, 89–97.

Draper, K., & Ponsford, J. (2008). Cognitive functioning ten years following traumatic brain injury and rehabilitation. Neuropsychology, 22(5), 618–625. doi:10.1037/0894-4105.22.5.618

Dunning, D. L., Westgate, B., & Adlam, A. L. (2016). A meta-analysis of working memory impairments in survivors of moderate-to-severe traumatic brain injury. Neuropsychology, 30(7), 811–819. doi:10.1037/neu0000285

Emch, M., von Bastian, C. C., & Koch, K. (2019). Neural Correlates of Verbal Working Memory: An fMRI Meta-Analysis. Front Hum Neurosci, 13, 180. doi:10.3389/fnhum.2019.00180

Felmingham, K. L., Baguley, I. J., & Green, A. M. (2004). Effects of diffuse axonal injury on speed of information processing following severe traumatic brain injury. Neuropsychology, 18(3), 564–571. doi:10.1037/0894-4105.18.3.564

Fitzpatrick, C., Archambault, I., Janosz, M., & Pagani, L. S. (2015). Early childhood working memory forecasts high school dropout risk. Intelligence, 53, 160–165.

Fletcher, P. C., & Henson, R. N. A. (2001). Frontal lobes and human memory: insights from functional neuroimaging. Brain, 124(5), 849–881.

Geday, J., Gjedde, A., Boldsen, A.-S., & Kupers, R. (2003). Emotional valence modulates activity in the posterior fusiform gyrus and inferior medial prefrontal cortex in social perception. Neuroimage, 18(3), 675–684.

Guerrero-Pedraza, A., McKenna, P., Gomar, J., Sarro, S., Salvador, R., Amann, B., … Pomarol-Clotet, E. (2012). First-episode psychosis is characterized by failure of deactivation but not by hypo-or hyperfrontality. Psychological medicine, 42(1), 73.

Hayter, A., Langdon, D., & Ramnani, N. (2007). Cerebellar contributions to working memory. Neuroimage, 36(3), 943–954.

Hillary, F. G. (2008). Neuroimaging of working memory dysfunction and the dilemma with brain reorganization hypotheses. J Int Neuropsychol Soc, 14(4), 526–534. doi:10.1017/S1355617708080788

Jacques, C., Jonas, J., Maillard, L., Colnat-Coulbois, S., Koessler, L., & Rossion, B. (2019). The inferior occipital gyrus is a major cortical source of the face-evoked N170: Evidence from simultaneous scalp and intracerebral human recordings. Human brain mapping, 40(5), 1403–1418.

Jaeggi, S. M., Buschkuehl, M., Perrig, W. J., & Meier, B. (2010). The concurrent validity of the N-back task as a working memory measure. Memory, 18(4), 394–412.

Jensen, O., Gelfand, J., Kounios, J., & Lisman, J. E. (2002). Oscillations in the alpha band (9–12 Hz) increase with memory load during retention in a short-term memory task. Cerebral cortex, 12(8), 877–882.

Kasahara, M., Menon, D. K., Salmond, C. H., Outtrim, J. G., Tavares, J. V., Carpenter, T. A., … Stamatakis, E. A. (2011). Traumatic brain injury alters the functional brain network mediating working memory. Brain Inj, 25(12), 1170–1187. doi:10.3109/02699052.2011.608210

Kim, C., Kroger, J. K., Calhoun, V. D., & Clark, V. P. (2015). The role of the frontopolar cortex in manipulation of integrated information in working memory. Neuroscience letters, 595, 25–29.

Kim, H. (2019). Neural activity during working memory encoding, maintenance, and retrieval: A network-based model and meta-analysis. Human brain mapping, 40(17), 4912–4933.

Klingberg, T., Forssberg, H., & Westerberg, H. (2002). Increased brain activity in frontal and parietal cortex underlies the development of visuospatial working memory capacity during childhood. Journal of Cognitive Neuroscience, 14(1), 1–10.

Lee, A. C., & Rudebeck, S. R. (2010). Investigating the interaction between spatial perception and working memory in the human medial temporal lobe. Journal of Cognitive Neuroscience, 22(12), 2823–2835.

Mak, L. E., Minuzzi, L., MacQueen, G., Hall, G., Kennedy, S. H., & Milev, R. (2017). The default mode network in healthy individuals: a systematic review and meta-analysis. Brain connectivity, 7(1), 25–33.

Mattay, V. S., Fera, F., Tessitore, A., Hariri, A. R., Berman, K. F., Das, S., … Weinberger, D. R. (2006). Neurophysiological correlates of age-related changes in working memory capacity. Neuroscience letters, 392(1-2), 32-37.

Mayer, J. S., Roebroeck, A., Maurer, K., & Linden, D. E. (2010). Specialization in the default mode: Task-induced brain deactivations dissociate between visual working memory and attention. Human brain mapping, 31(1), 126–139.

McAllister, T. W., Sparling, M. B., Flashman, L. A., Guerin, S. J., Mamourian, A. C., & Saykin, A. J. (2001). Differential working memory load effects after mild traumatic brain injury. Neuroimage, 14(5), 1004–1012. doi:10.1006/nimg.2001.0899

McGeown, W. J., Mazzoni, G., Venneri, A., & Kirsch, I. (2009). Hypnotic induction decreases anterior default mode activity. Consciousness and cognition, 18(4), 848–855.

McKiernan, K. A., Kaufman, J. N., Kucera-Thompson, J., & Binder, J. R. (2003). A parametric manipulation of factors affecting task-induced deactivation in functional neuroimaging. Journal of Cognitive Neuroscience, 15(3), 394–408.

McVay, J. C., & Kane, M. J. (2012). Why does working memory capacity predict variation in reading comprehension? On the influence of mind wandering and executive attention. Journal of experimental psychology: general, 141(2), 302.

Migo, E., Mitterschiffthaler, M., O’Daly, O., Dawson, G., Dourish, C., Craig, K., … Jackson, S. (2015). Alterations in working memory networks in amnestic mild cognitive impairment. Aging, Neuropsychology, and Cognition, 22(1), 106–127.

Murty, V. P., Sambataro, F., Radulescu, E., Altamura, M., Iudicello, J., Zoltick, B., … Mattay, V. S. (2011). Selective updating of working memory content modulates meso-cortico-striatal activity. Neuroimage, 57(3), 1264–1272.

Muschelli, J., Nebel, M. B., Caffo, B. S., Barber, A. D., Pekar, J. J., & Mostofsky, S. H. (2014). Reduction of motion-related artifacts in resting state fMRI using aCompCor. Neuroimage, 96, 22–35. doi:10.1016/j.neuroimage.2014.03.028

Narayanan, N. S., Prabhakaran, V., Bunge, S. A., Christoff, K., Fine, E. M., & Gabrieli, J. D. (2005). The role of the prefrontal cortex in the maintenance of verbal working memory: an event-related FMRI analysis. Neuropsychology, 19(2), 223.

Newsome, M. R., Steinberg, J. L., Scheibel, R. S., Troyanskaya, M., Chu, Z., Hanten, G., … Levin, H. S. (2008). Effects of traumatic brain injury on working memory-related brain activation in adolescents. Neuropsychology, 22(4), 419–425. doi:10.1037/0894-4105.22.4.419

Owen, A. M., McMillan, K. M., Laird, A. R., & Bullmore, E. (2005). N-back working memory paradigm: A meta-analysis of normative functional neuroimaging studies. Human brain mapping, 25(1), 46–59.

Perlstein, W. M., Cole, M. A., Demery, J. A., Seignourel, P. J., Dixit, N. K., Larson, M. J., & Briggs, R. W. (2004). Parametric manipulation of working memory load in traumatic brain injury: behavioral and neural correlates. J Int Neuropsychol Soc, 10(5), 724–741. doi:10.1017/S1355617704105110

Pomarol-Clotet, E., Salvador, R., Sarro, S., Gomar, J., Vila, F., Martinez, A., … Capdevila, A. (2008). Failure to deactivate in the prefrontal cortex in schizophrenia: dysfunction of the default mode network? Psychological medicine, 38(8), 1185.

Ponsford, J., Downing, M. G., Olver, J., Ponsford, M., Acher, R., Carty, M., & Spitz, G. (2014). Longitudinal follow-up of patients with traumatic brain injury: outcome at two, five, and ten years post-injury. J Neurotrauma, 31(1), 64–77. doi:10.1089/neu.2013.2997

Ponsford, J., Sloan, S., & Snow, P. (2012). Traumatic Brain Injury: Rehabilitation for Everyday Adaptive Living, 2nd Edition: Taylor & Francis.

Pyka, M., Beckmann, C. F., Schöning, S., Hauke, S., Heider, D., Kugel, H., … Konrad, C. (2009). Impact of working memory load on FMRI resting state pattern in subsequent resting phases. PLoS One, 4(9), e7198.

Raichle, M. E., MacLeod, A. M., Snyder, A. Z., Powers, W. J., Gusnard, D. A., & Shulman, G. L. (2001). A default mode of brain function. Proceedings of the National Academy of Sciences, 98(2), 676–682.

Ramnani, N. (2006). The primate cortico-cerebellar system: anatomy and function. Nature Reviews Neuroscience, 7(7), 511–522.

Rossi, A. F., Pessoa, L., Desimone, R., & Ungerleider, L. G. (2009). The prefrontal cortex and the executive control of attention. Experimental brain research, 192(3), 489.

Rottschy, C., Langner, R., Dogan, I., Reetz, K., Laird, A. R., Schulz, J. B., … Eickhoff, S. B. (2012). Modelling neural correlates of working memory: a coordinate-based meta-analysis. Neuroimage, 60(1), 830–846. doi:10.1016/j.neuroimage.2011.11.050

Sambataro, F., Murty, V. P., Callicott, J. H., Tan, H.-Y., Das, S., Weinberger, D. R., & Mattay, V. S. (2010). Age-related alterations in default mode network: impact on working memory performance. Neurobiology of aging, 31(5), 839–852.

Sanchez-Carrion, R., Fernandez-Espejo, D., Junque, C., Falcon, C., Bargallo, N., Roig, T.,… Vendrell, P. (2008). A longitudinal fMRI study of working memory in severe TBI patients with diffuse axonal injury. Neuroimage, 43(3), 421–429. doi:10.1016/j.neuroimage.2008.08.003

Sanchez-Carrion, R., Gomez, P. V., Junque, C., Fernandez-Espejo, D., Falcon, C., Bargallo, N., … Bernabeu, M. (2008). Frontal hypoactivation on functional magnetic resonance imaging in working memory after severe diffuse traumatic brain injury. J Neurotrauma, 25(5), 479–494. doi:10.1089/neu.2007.0417

Scherf, K. S., Sweeney, J. A., & Luna, B. (2006). Brain basis of developmental change in visuospatial working memory. Journal of Cognitive Neuroscience, 18(7), 1045–1058.

Schon, K., Quiroz, Y. T., Hasselmo, M. E., & Stern, C. E. (2009). Greater working memory load results in greater medial temporal activity at retrieval. Cereb Cortex, 19(11), 2561–2571. doi:10.1093/cercor/bhp006

Sternberg, S. (1966). High-speed scanning in human memory. Science, 153(3736), 652–654.

Tallon-Baudry, C., Bertrand, O., & Fischer, C. (2001). Oscillatory synchrony between human extrastriate areas during visual short-term memory maintenance. Journal of Neuroscience, 21(20), RC177–RC177.

Tomasi, D., Ernst, T., Caparelli, E. C., & Chang, L. (2006). Common deactivation patterns during working memory and visual attention tasks: An intra-subject fMRI study at 4 Tesla. Human brain mapping, 27(8), 694–705.

Tomlinson, S. P., Davis, N. J., Morgan, H. M., & Bracewell, R. M. (2014). Cerebellar contributions to verbal working memory. The Cerebellum, 13(3), 354–361.

Tun, P. A., & Lachman, M. E. (2008). Age differences in reaction time and attention in a national telephone sample of adults: education, sex, and task complexity matter. Developmental psychology, 44(5), 1421.

