## Supplementary for "Aberrant modulation of brain activity underlies impaired working memory following traumatic brain injury"

Supplementary Materials

**Supplementary Table 1.** Clinical characteristics of TBI participants

| **Age (years)** | **Sex** | **Cause**  **of injury** | **PTA (days)^a^** | **GCS (lowest)^b^** | **Time since injury (months)** | **CT/MRI finding** |
| --- | --- | --- | --- | --- | --- | --- |
| 41 | F | Car accident | 19 | 10 | 0.8 | NAD |
| 38 | F | Motorcycle | 7 | 3 | 1 | SAH, SDH |
| 62 | M | Fall of mower | 7 | 14 | 1.2 | Contusion, EDH, SAH |
| 32 | M | Car accident | 22 | 12 | 6 | Contusion, ICH, SAH |
| 48 | M | Motorcycle | 5 | 14 | 3.9 | NAD |
| 45 | M | Bicycle vs. Car | 21 | 8 | 0.7 | SAH |
| 33 | F | Fall of bicycle | 22 | 6 | 1 | SAD |
| 52 | M | Car accident | 14 | 8 | 1.4 | Contusion |
| 21 | F | Pedestrian vs. Car | 3 | 13 | 1.4 | EDH, ICH, SAD |
| 19 | F | Car accident | 63 | 3 | 6.6 | Contusion, EDH, ICH, SAD |
| 22 | M | Car accident | 43 | 15 | 1.8 | SDH |
| 19 | M | Car accident | 24 | 3 | 1.1 | DAI, ICH |
| 19 | M | Car accident | 50 | 3 | 2.7 | DAI, ICH |
| 73 | F | Car accident | 22 | 4 | 2.6 | EDH, SAH, SDH |
| 56 | M | Motorcycle | 23 | 10 | 2.1 | ICH, SAH, SAH |
| 65 | M | Car accident | 24 | 10 | 1.2 | DAI, ICH |
| 32 | F | Car accident | 18 | 11 | 2.3 | ICH |
| 27 | F | Car accident | 7 | 14 | 3.1 | SAH |
| 24 | M | Motorcycle | 39 | 3 | 1.6 | DAI, IVH, SAH |
| 42 | M | Bicycle vs. Truck | 8 | 14 | 1.1 | ICH, SAH, SDH |
| 18 | M | Car accident | 28 | 6 | 1.4 | Contusion, SAH, SDH |
| 18 | M | Pedestrian vs. Bus | 22 | 7 | 1.8 | ICH |
| 62 | M | Motorcycle | Unknown | 15 | 3.4 | SHD |
| 50 | M | Motorcycle | 18 | 13 | 1.2 | ICH |
| 41 | M | Pedestrian vs. Tram | 30 | 6 | 2.5 | ICH |
| 67 | M | Bicycle vs. car | 9 | 8 | 26.93 | DAI, ICH |
| 25 | F | Fall from horse | 34 | 6 | 34.82 | ICH |
| 67 | M | Car accident | 7 | 10 | 19.33 | ICH |
| 42 | M | Car accident | 180 | 4 | 28.34 | DAI, SAH |
| 46 | M | Motorcycle | 15 | 13 | 23.31 | DAI |
| 46 | M | Pedestrian vs. car | 33 | 15 | 13.35 | NAD |
| 50 | M | Motorcycle | 43 | 6 | 26.17 | Contusion, DAI, ICH |
| 62 | M | Car accident | Unknown | Unknown | 28.57 | Pneumocephalus |
| 44 | M | Motorcycle | 34 | 8 | 15.75 | ICH, pneumocephalus, SAH |
| 61 | M | Motorcycle | 23 | 14 | 15.42 | NAD |
| 23 | M | Car accident | 14 | 13 | 17.19 | Contusion, ICH, SAH |
| 20 | F | Pedestrian vs. car | 41 | 3 | 27.58 | Contusion, SAH, SDH |
| 64 | M | Motorcycle | 46 | 9 | 33.70 | DAI, ICH |
| 28 | F | Car accident | 11 | ? | 17.42 | ICH |
| 27 | F | Bicycle vs. tram | 21 | 13 | 21.17 | Contusion, SDH |
| 31 | M | Motorcycle | 24 | 14 | 34.82 | ICH |
| 49 | M | Pedestrian vs. car | 10 | 8 | 16.73 | ICH, SDH |
| 42 | M | Car accident | 18 | 3 | 21.34 | ICH, SAH, SDH |

GSC = Glasgow Coma Scale; PTA = post-traumatic amnesia; NAD = no abnormality detected; SAD = subarachnoid haemorrhage, SHD = subdural haemorrhage, EDH = extradural haematoma; ICH = intracerebral haemorrhage, DAI = diffuse axonal injury.

^a^ PTA duration were available for *n* = 41 TBI participants.

^b^ Acute GCS were available for *n* = 42 TBI participants.

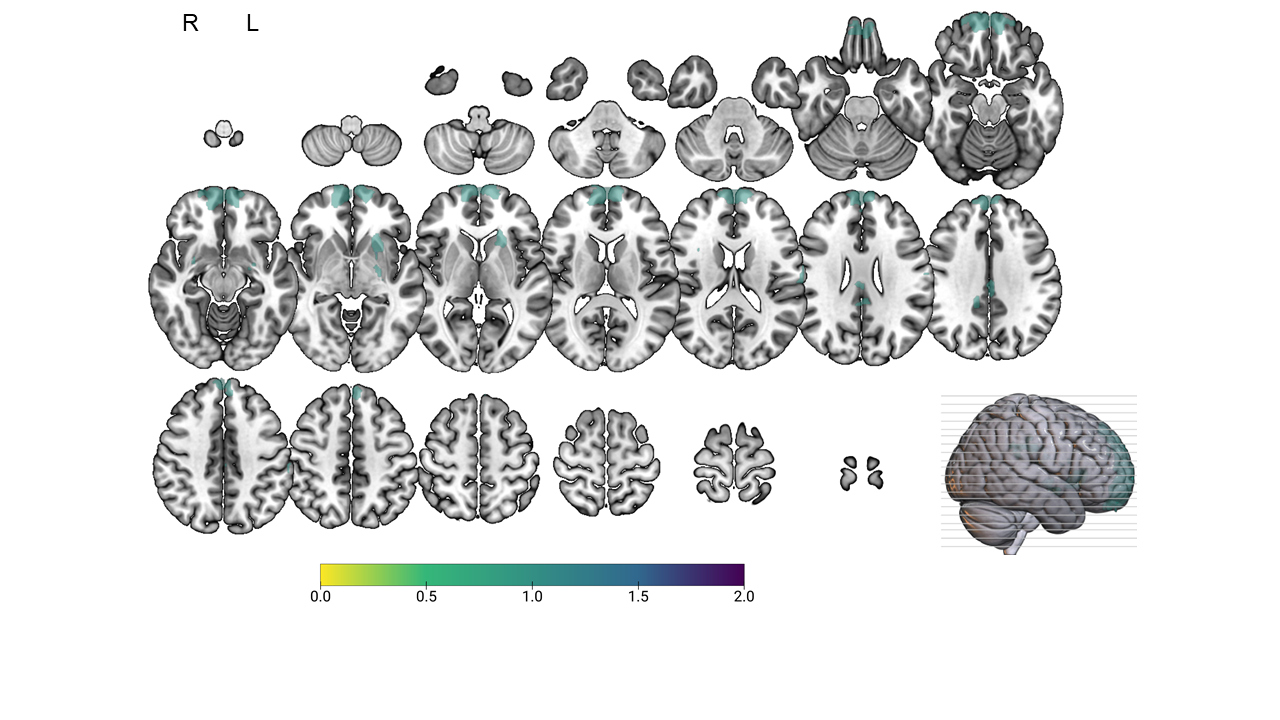

**Supplementary Figure 1.** Lesion overlay map that characterise brain pathology of TBI participants in MNI space. Pathology was predominantly evident in frontal, temporal, and subcortical areas. Note: purple colour indicates greater lesion overlap across participants.

**Detailed MRI Preprocessing**

Preprocessing of MRI data involved the following: each T1-weighted (T1w) image was corrected for intensity non-uniformity (INU) with N4BiasFieldCorrection (Tustison et al., 2010) and skull-stripped with a Nipype implementation of the antsBrainExtraction.sh workflow (using OASIS30ANTs as target template). Brain tissue segmentation of cerebrospinal fluid (CSF), white-matter (WM) and grey-matter (GM) was performed on the brain-extracted T1w using FAST (FSL 5.0.9; Zhang, Brady, & Smith, 2001). Cortical brain surfaces were reconstructed using recon-all (FreeSurfer 6.0.1; Dale, Fischl, & Sereno, 1999). Volume-based spatial normalisation to two standard spaces (MNI152NLin6Asym, MNI152NLin2009cAsym) was performed through nonlinear registration with antsRegistration (ANTs 2.2.0).

Functional data were skull-stripped using a custom methodology of fMRIPrep. fMRIPrep’s fieldmap-less approach was used to correct for susceptibility distortion using a deformation field resulting from co-registering the BOLD reference to the same-subject T1w-reference with its intensity inverted (Huntenburg, 2014; Wang et al., 2017).

The BOLD reference was then co-registered to the T1w reference using boundary-based registration (Greve & Fischl, 2009) with six degrees of freedom with bbregister (FreeSurfer). Head-motion parameters with respect to the BOLD reference (transformation matrices, and six corresponding rotation and translation parameters) were estimated using FSL’s MCFLIRT (Jenkinson, Bannister, Brady, & Smith, 2002). The BOLD time-series were resampled onto their original, native space by applying a single, composite transform to correct for head-motion and susceptibility distortions. The BOLD time-series were resampled into MNI152NLin6Asym standard space.

Non-steady state volumes were removed from preprocessed BOLD images and spatial smoothing with an isotropic, Gaussian kernel of 6mm FWHM (full-width half-maximum) was applied. Several confounding time-series were calculated based on the preprocessed BOLD: framewise displacement (FD), DVARS and three region-wise global signals. FD and DVARS are calculated, both using their implementations in Nipype (Power et al., 2014). The three global signals are extracted within the CSF, the WM, and the whole-brain masks. Additionally, a set of physiological regressors were extracted to allow for component-based noise correction (CompCor; Behzadi, Restom, Liau, & Liu, 2007). Principal components are estimated after high-pass filtering the preprocessed BOLD time-series (using a discrete cosine filter with 128s cut-off) for the two CompCor variants: temporal (tCompCor) and anatomical (aCompCor). tCompCor components are then calculated from the top 5% variable voxels within a mask covering the subcortical regions. For aCompCor, components are calculated within the intersection of the aforementioned mask and the union of CSF and WM masks calculated in T1w space, after their projection to the native space of each functional run (using the inverse BOLD-to-T1w transformation). Components are also calculated separately within the WM and CSF masks.

Many internal operations of fMRIPrep use Nilearn 0.6.1 (Abraham et al., 2014), mostly within the functional processing workflow. For more details of the pipeline, see <https://fmriprep.readthedocs.io/en/1.0.8/workflows.html>.

**Supplementary Table 2.** Significant clusters during the Sternberg working memory task for all participants

| **Contrast** | **Region** | **Voxels** | **Peak coordinates** | | | ***P*-value** |
| --- | --- | --- | --- | --- | --- | --- |
|  |  |  | X | Y | Z |  |
| Encoding | Occipital_Mid_R  Occipital_Mid_L  Precentral_L  Occipital_Mid_R  Precentral_R  Supp_Motor_Area_L  Cerebellum_8_R  Insula_L  Cerebellum_8_L  Insula_R  Temporal_Mid_L  Temporal_Sup_R  Postcentral_L  Cingulum_Mid_L  Vermis_9 | 4572  4389  3567  1700  1623  1398  467  447  336  247  246  205  127  91  83 | 28  -36  -52  32  52  -6  28  -32  -28  30  -52  48  -54  -6  0 | -90  -84  0  -80  12  2  -64  16  -66  28  -48  -40  -18  -20  -54 | 2  2  46  36  36  60  -54  10  -52  -2  2  14  18  48  -36 | <0.001  <0.001  <0.001  <0.001  <0.001  <0.001  <0.001  <0.001  <0.001  <0.001  <0.001  <0.001  <0.001  0.007  0.013 |
| Maintenance | Calcarine_L  Cerebellum_Crus1_R  Precentral_L  Supp_Motor_Area_L  Insula_L  Cerebellum_8_R  Cerebellum_Crus1_L  Rolandic_Oper_L  Insula_R | 656  416  407  371  324  255  171  84  64 | 0  42  -38  -6  -28  38  -40  -46  32 | -86  -66  -14  8  26  -58  -62  0  26 | -8  -28  68  56  2  -54  -32  12  4 | <0.001  <0.001  <0.001  <0.001  <0.001  <0.001  <0.001  0.013  0.049 |
| Retrieval | Frontal_Inf_Oper_L  Supp_Motor_Area_L  Insula_R  Insula_L  Parietal_Inf_L  Occipital_Inf_L  Calcarine_R  Precentral_R  Occipital_Inf_R  Calcarine_L  Frontal_Mid_L  Cerebellum_6_R  Lingual_R | 2370  1920  928  697  562  363  202  195  164  95  86  64  63 | -36  -6  32  -30  -28  -28  14  44  30  -14  -26  20  38 | 8  16  26  24  -54  -94  -78  6  -96  -70  44  -52  -84 | 30  46  2  6  42  -10  10  30  0  8  26  -20  -16 | <0.001  <0.001  <0.001  <0.001  <0.001  <0.001  <0.001  <0.001  <0.001  0.006  0.011  0.046  0.049 |

**Supplementary Table 3.** Significant clusters overall (collapsed across the various load and stage)

| **Contrast** | **Region** | **Voxels** | **Peak coordinates** | | | ***P*-value** |
| --- | --- | --- | --- | --- | --- | --- |
|  |  |  | X | Y | Z |  |
| TBI < HC | Cerebellum_Crus1_L  Frontal_Mid_L  Temporal_Sup_L | 90  77  70 | -40  -32  -56 | -70  40  -42 | -18  38  22 | 0.014  0.031  0.048 |

**Supplementary Table 4.** Significant clusters during the low cognitive load condition

| **Contrast** | **Region** | **Voxels** | **Peak coordinates** | | | ***P*-value** |
| --- | --- | --- | --- | --- | --- | --- |
|  |  |  | X | Y | Z |  |
| TBI < HC | Cerebellum_Crus1_L  Frontal_Mid_R  Temporal_Sup_L  Frontal_Sup_L | 115  100  70  70 | -40  38  -60  -26 | -66  50  -38  38 | -24  32  22  38 | 0.003  0.007  0.042  0.042 |

**Supplementary Table 5.** Significant clusters during the high > low load condition

| **Contrast** | **Region** | **Voxels** | **Peak coordinates** | | | ***P*-value** |
| --- | --- | --- | --- | --- | --- | --- |
|  |  |  | X | Y | Z |  |
| TBI > HC | Parietal_Sup_R  Temporal_Inf_R  Frontal_Mid_R  Occipital_Inf_L | 126  109  72  66 | 36  50  32  -50 | -52  -54  14  -72 | 58  -16  64  -14 | <0.001  0.002  0.019  0.030 |

**Supplementary Table 6.** Significant clusters during maintenance

| **Contrast** | **Region** | **Voxels** | **Peak coordinates** | | | ***P*-value** |
| --- | --- | --- | --- | --- | --- | --- |
|  |  |  | X | Y | Z |  |
| TBI < HC | Frontal_Sup_R | 96 | 18 | 60 | 18 | 0.006 |

**Supplementary Table 7.** Significant clusters during encoding high > low

| **Contrast** | **Region** | **Voxels** | **Peak coordinates** | | | ***P*-value** |
| --- | --- | --- | --- | --- | --- | --- |
|  |  |  | X | Y | Z |  |
| TBI > HC | Occipital_Inf_L | 73 | -46 | -78 | -12 | 0.018 |

**Supplementary Table 8.** Significant clusters during maintenance high > low

| **Contrast** | **Region** | **Voxels** | **Peak coordinates** | | | ***P*-value** |
| --- | --- | --- | --- | --- | --- | --- |
|  |  |  | X | Y | Z |  |
| TBI < HC | Cerebellum_Crus1_L  Calcarine_L  Cerebellum_Crus1_R | 108  75  62 | -44  -8  44 | -64  -102  -68 | -20  -10  -20 | 0.002  0.015  0.039 |

Activity during encoding is associated with behaviour

In the main text, we found several brain regions presenting with aberrant neural processing following TBI. Aberrant regional activity was present only under specific working memory conditions. For example, when all trials were averaged, with increased load demands, or only for certain working memory subcomponents. We further probed these regions to examine whether activity in specific brain regions was directly related to working memory capacity. To do this, we extracted each individual’s COPE statistic and correlated this with working memory accuracy (using Spearman correlations). We found that a greater reduction in activity in the left inferior occipital gyrus with *greater* task demands was associated with better task accuracy. More specifically, there was a significant negative correlation between COPE values and low (*r*_s_ = -0.381, *P =* 0.023; Supplementary Fig. 2A) and high dprime scores (*r*_s_ = -0.416, *P =* 0.013; Supplementary Fig. 2B). To investigate whether these association differed for each group, we conducted separate correlations for the TBI group and healthy controls. Although there was no significant correlation between COPE values in the low load condition and low dprime scores for the TBI group (*r*_s_ = -0.050, *P =* 0.844; Supplementary Fig. 2C), a significant negative correlation emerged for healthy controls (*r*_s_ = -0.636, *P =* 0.006; Supplementary Fig. 2D). Further analysis indicated that this pattern of finding was specifically driven by activity in the low load condition. That is, there was a significant correlation between COPE values in the low load condition and low dprime scores (*r*_s_ = 0.569, *P =* 0.017; Supplementary Fig. 2E). In contrast, there was no correlation between COPE values in the high load condition and high dprime scores (*r*_s_ = 0.184, *P =* 0.480; Supplementary Fig. 2F). Taken together, results suggest that greater deactivation as task demands increased is associated with better working memory accuracy in the low condition.

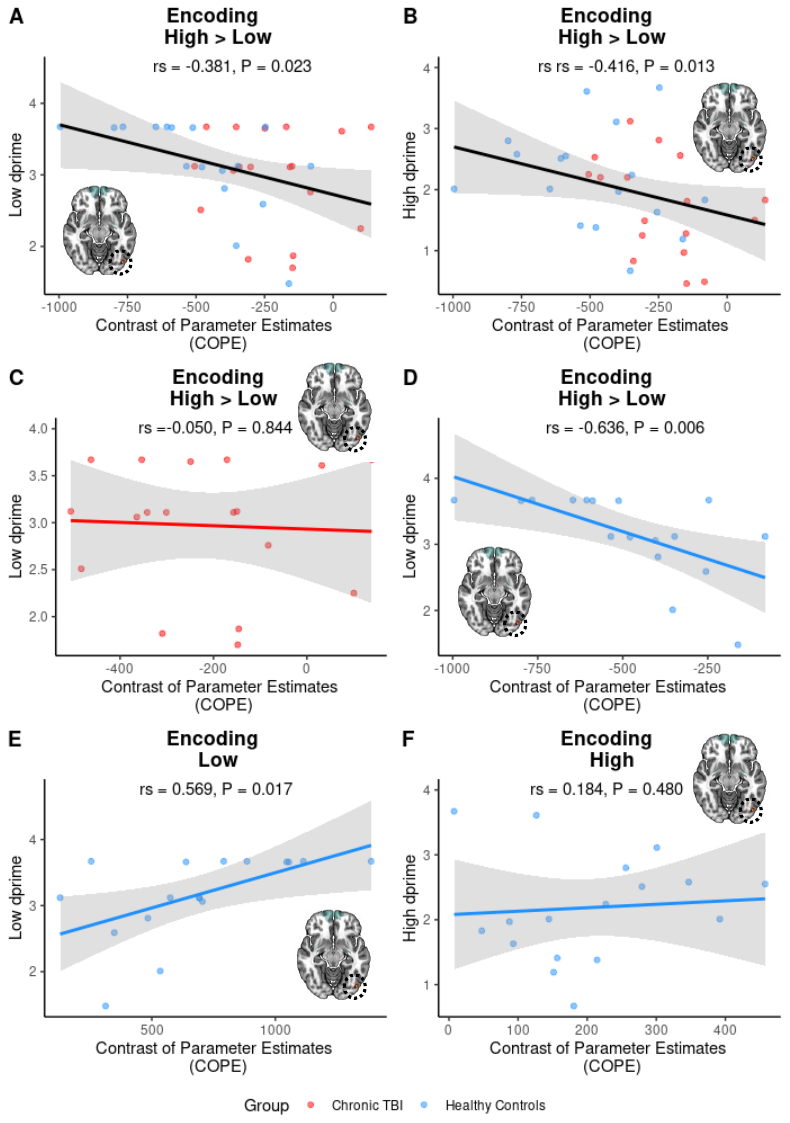

**Supplementary Figure 2.** Increased activation in the left inferior occipital gyrus during encoding was associated with better working memory accuracy in the low load condition. A) Overall, there was a significant negative correlation between COPE values associated with increased cognitive load and low dprime scores. B) There was also a significant correlation between COPE values associated with increased cognitive load and high dprime scores. C) Additional analysis indicated that the correlation between COPE values and low dprime scores was not significant for the TBI group. D) There was, however, a significant correlation between COPE values and low dprime scores for healthy controls. E) Further investigation indicated that the association was specifically driven by activity in the low load condition. That is, there was a significant correlation between COPE values in the low load condition and low dprime scores. F) In contrast, there was no significant correlation between COPE values in the high load condition and high dprime scores.
